# Reduced glutathione levels in *Enterococcus faecalis* trigger metabolic and transcriptional compensatory adjustments during iron exposure

**DOI:** 10.1101/2025.04.04.647250

**Authors:** Víctor Aliaga-Tobar, Jorge Torres, Sebastián Mendoza, Gabriel Gálvez, Jaime Ortega, Sebastián Gómez, Valentina Parra, Felipe Arenas, Alejandro Maass, Anne Siegel, Mauricio González, Mauricio Latorre

## Abstract

*Enterococcus faecalis*, a facultative anaerobic pathogen and common constituent of the gastrointestinal microbiota, must navigate varying iron levels within the host. This study explores its response to iron supplementation in a glutathione-deficient mutant strain (Δ*gsh*). We examined the transcriptomic and metabolic responses of a glutathione synthetase mutant strain (Δ*gsh*) exposed to iron supplementation, integrating these data into a genome-scale metabolic model (GSMM). Our results show that under glutathione deficiency, *E. faecalis* reduces intracellular iron levels and shifts its transcriptional response to prioritize energy production genes. Notably, basal metabolites, including arginine, increase. The GSMM highlights the importance of arginine metabolism, particularly the *arc* operon (anaerobic arginine catabolism), as a compensatory mechanism for reduced glutathione during iron exposure. These findings provide insights into how *E. faecalis* adjusts metal homeostasis and transcriptional/metabolic processes to mitigate the effects of oxidative stress caused by iron.

**IMPORTANCE:** Iron is essential for bacterial survival, yet its excess can be harmful through increase of oxidative stress. *Enterococcus faecalis*, a bacterium common member of the human gut, must carefully balance its iron levels in order to survive in changing environments. This study studies how *E. faecalis* compensates the reduced levels of glutathione —a key antioxidant— when exposed to high iron concentrations. We discovered that *E. faecalis* lowers its intracellular iron levels under glutathione decrease and reprograms its metabolism to prioritize energy production. These findings provide valuable insights into bacterial adaptation mechanisms under oxidative stress conditions, which could influence the development of new strategies to combat bacterial infections.

## INTRODUCTION

Since the emergence of oxygen on Earth, cells have faced the dual-edged nature of O₂. On one hand, oxygen serves as the terminal electron acceptor in a series of intricate redox reactions, such as oxidative phosphorylation. On the other hand, it is a stable allotropic molecule with an electronic configuration containing two unpaired electrons, capable of generating reactive oxygen species (ROS) (1, 2). These ROS are particularly toxic to cells due to their high reactivity, causing damage to cellular components, including DNA, membrane lipids, and proteins—a process known as oxidative stress, which can ultimately lead to cell death (3). Given the omnipresence of oxidative stress in biological systems, bacteria have evolved sophisticated mechanisms to mitigate its harmful effects, particularly in metal-rich environments. However, the precise role of key antioxidant molecules in modulating global transcriptional responses under such conditions remains underexplored.

In bacterial species growing under aerobic conditions, a portion of the cellular damage caused by oxidative stress can be attributed to ROS formed endogenously through reactions between O₂ and univalent electron donors, such as metals (3, 4). Among these metals, iron is particularly harmful. While iron is essential for many bacterial species due to its redox activity (5), it becomes toxic at elevated concentrations. Through Fenton and/or Haber-Weiss reactions, iron reacts with hydrogen peroxide (H₂O₂) and superoxide (O₂•−), generating hydroxyl radicals (OH•), which can damage various biological macromolecules (5, 6). Additionally, prolonged oxidative stress can trigger the release of iron atoms from mononuclear enzymes, further increasing intracellular iron levels and exacerbating cellular damage (7). Thus, tight regulation of both oxidative stress and iron homeostasis is critical for bacterial survival.

Regarding antioxidative defense, these systems comprise both non-enzymatic and enzymatic components, along with the regulatory networks that govern them (1, 8). Specifically, enzymatic components include enzymes targeting specific ROS, while non-enzymatic modules involve small molecules with antioxidant potential (9), among which the tripeptide glutathione is the most well-characterized example (1, 10, 11).

Glutathione is the most abundant antioxidant molecule in cells and is present in virtually all bacteria and eukaryotic cells (4). It plays a crucial role not only in protecting against oxidative stress but also in maintaining cellular homeostasis, regulating sulfur transport, conjugating metabolites, detoxifying xenobiotics, conferring antibiotic resistance, regulating enzyme activity, and modulating the expression of stress response genes (11). Glutathione protects against oxidative stress through direct and indirect interactions with ROS.

In these mechanisms, glutathione donates electrons directly to O₂•−, OH•, peroxy radicals (ROO•), and peroxynitrite (ONOO−), converting glutathione to glutathione disulfide (GSSG). Additionally, glutathione peroxidase decomposes H₂O₂ using glutathione (4). Finally, glutathione reductase regenerates glutathione from GSSG, allowing for the recycling of this antioxidant during redox processes (4). Beyond its role in redox homeostasis, recent studies suggest that glutathione also plays an integral part in metal tolerance.

For instance, in *Streptococcus pyogenes*, glutathione has been shown to buffer excess copper under saturation conditions, conferring additional tolerance and complementing efflux systems to maintain metal homeostasis (12). While these findings highlight the relevance of glutathione in bacterial stress responses, little is known about how glutathione depletion impacts the global transcriptional landscape of bacteria, particularly under conditions of iron-induced oxidative stress.

In recent years, *Enterococcus faecalis* has emerged as a well-established model for studying systems biology and heavy metal stress responses (6, 13–17). Faced with an excess of Fe, this bacterium activates or represses about 15% of its total genes, classifying this response to the metal as global, here genes directly involved in iron homeostasis proteins as well as ROS resistance are prominently regulated (6, 15).

Regarding *E. faecalis* and the role of glutathione as an antioxidant, it has been observed that in oxidative environments, the levels of this antioxidant increase (18). Additionally, the enzyme glutathione reductase, which is responsible for maintaining the supply of reduced glutathione by converting GSSG to GSH, has been purified and characterized (19). However, despite its well-documented role in oxidative stress defense, the mechanisms governing glutathione biosynthesis in *E. faecalis* remain largely unknown.

To address this gap, we applied an integrative systems biology approach to investigate the global transcriptional and metabolic responses of the pathogenic bacterium *E. faecalis* under elevated iron levels in a glutathione-deficient scenario. To our knowledge, this is the first report that studies at a systemic level, the importance of glutathione in dealing with iron induced oxidative stress, raising the question of whether this reduction in glutathione content could also cause global metabolic adjustments that allow the bacterium to cope with the stress scenario.

## RESULTS

### 1. Glutathione synthetase of *E. faecalis* is conserved in *Lactobacillales*

Typically, the synthesis of glutathione involves the sequential action of two enzymes: γ-glutamylcysteine synthetase (γGCS) and glutathione synthetase (GS). However, in some species, such as *Streptococcus agalactiae*, a member of the *Lactobacillale* order, glutathione synthesis is catalyzed by a single enzyme that combines both functions: γGCS-GS (20). In the *E. faecalis* genome, we identified a homologous γGCS-GS protein encoded by the monocistronic gene gshF-like (EF3089), which consists of an ORF of 2,271 nucleotides and encodes a 756-amino acid protein (**Figure 1A**).

**Figure 1.**
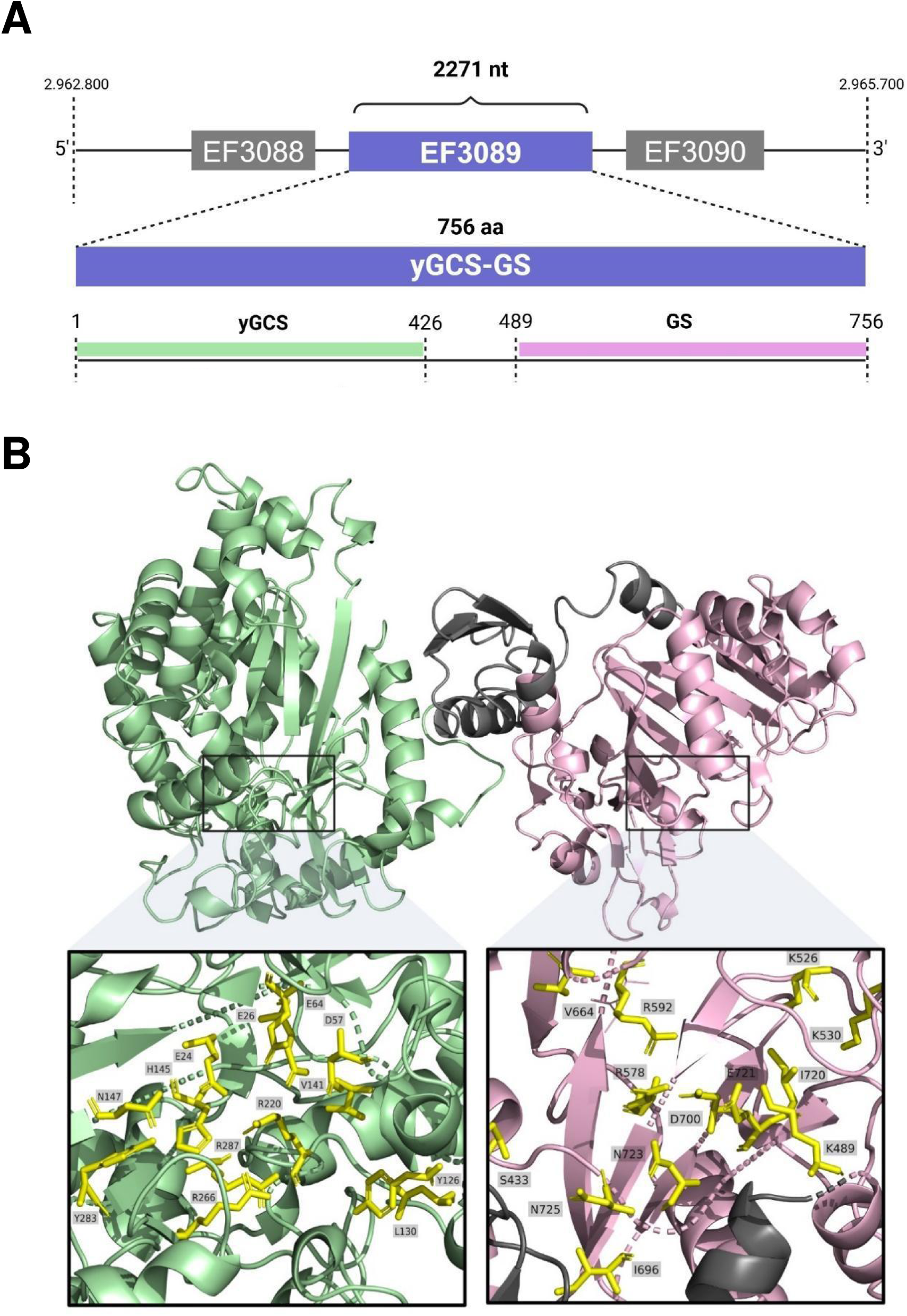
Schematic representation of the bifunctional γGCS-GS enzyme from *E. faecalis*. **A.** Genomic context model of gen EF3089, encoding the bifunctional enzyme γGCS-GS. **B.** Domain concatenation model of the γGCS-GS enzyme, with the γGCS and GS domains highlighted in green and pink, respectively. Key conserved residues in the γGCS and GS domains are depicted in yellow. The three-dimensional structure of γGCS-GS was reconstructed using the Alphafold protein structure database.

A comparative analysis of γGCS-GS across *Lactobacillales* species revealed a high degree of conservation in its primary structure (identity values > 45%) (**Sup.** Figure 1), suggesting a common evolutionary origin shared with *Lactobacillus plantarum* and *S. agalactiae* (20).

To validate the predicted gene annotation *in silico*, we constructed a homology-based model using the crystallized structure of γGCS-GS from *Streptococcus agalactiae* (48% identity and 66% similarity between both protein sequences) (21). The resulting structural model (**Figure 1B**) confirms a predominantly alpha-helical structure with approximately five beta-sheet domains. Additionally, the functional domain from 441 to 464, necessary for full γGCS activity, is well conserved in the *E. faecalis* enzyme.

The substrate-binding region (residues 448–489), crucial for GS function in *S. agalactiae*, is also conserved in *E. faecalis*, reinforcing its predicted enzymatic role (20).

Furthermore, it has been found that the γGCS-GS enzyme can be inhibited by its glutathione product (20). We identified that the inhibitory residues located between 495 and 508 in *S. agalactiae* are highly conserved in *E. faecalis*, further supporting the functional similarity between these orthologs.

Regarding specific amino acids, we identified key residues involved in four processes relevant to enzymatic action: cysteine binding, glutamate binding, glycine binding and magnesium cofactor binding. In *S. agalactiae*, the coordination of the magnesium cofactor involves three distinct sites, with residues Glu-29, Asp-60, Glu-67, His-150, Glu-328, and Glu-27.

Similarly, the essential residue for glycine binding is an arginine located at position 588. The glutamate binding residues are Ile-146 (there is in 145), Arg-235 (not present), His-150. Finally, the primary residues for cysteine binding in γGCS-GS—Phe-61, Tyr-131 (129 in *E. faecalis*), and Leu-135 (133 in *E. faecalis*)—are also present, albeit with minor positional shifts (20, 22, 23).

In *E. faecalis*, Arg-588 (glycine binding), Tyr-131, and Leu-135 are located at Arg-592, Tyr-129, and Leu-133, respectively. Although these residues do not align perfectly with their counterparts in *S. agalactiae*, their conservation suggests they likely retain similar functional properties, warranting further experimental validation.

Overall, the strong sequence conservation within *Lactobacillales* and the presence of well-preserved functional residues support the conclusion that the annotated gshF-like gene in *E. faecalis* encodes a functional γGCS-GS enzyme, reinforcing its likely role in glutathione biosynthesis and oxidative stress response.

### 2. Absence of γGCS-GS in *E. faecalis* induces the reduction of intracellular glutathione and iron content

The increase in intracellular iron content induces the production of ROS through a series of chemical reactions (24). It is known that *E. faecalis* has developed mechanisms to maintain homeostasis and respond to conditions of iron excess or deficiency (6, 15). These mechanisms include transcriptional changes that promotes the activation of genes and regulators related to basal metabolism, mainly controlled by Fur, a transcription factor responsible for regulating metal uptake systems to prevent excessive intracellular iron accumulation (6). However, little is known about the mechanisms involved in the control of iron-generated toxicity in *E. faecalis*.

Considering this, an initial question in our study was if absence of glutathione could be harmful to *E. faecalis* under conditions of iron excess. To explore this, we studied the mutant strain for the γGCS-GS enzyme (Δ*gsh*) and compared its viability in relation to the wild-type (WT) strain under different levels of iron exposure. Surprisingly, the results (**Figure 2A**) indicate no significant differences in viability between both strains, suggesting that the absence of the γGCS-GS enzyme does not impair growth, even under high iron exposure (4 mM, **Sup.** Figure 2). While the viability results did not show impaired growth in both WT and Δ*gsh* strains, given the importance of glutathione as an antioxidant, it is plausible to assume some compensatory effect that helps reduce iron toxicity.

**Figure 2.**
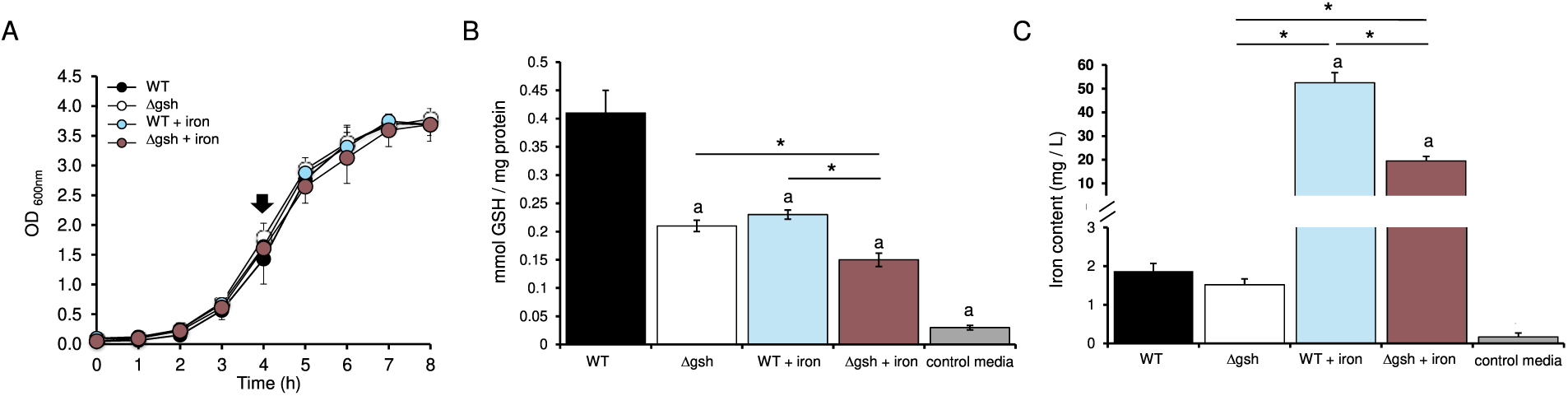
Metal homeostatic response of *E. faecalis* WT and *Δgsh* strains under iron excess. The iron exposure condition corresponds to 0.5 mM of FeCl_3_ per 3 h **A.** Growth curves. Black arrow indicates the point selected for quantification of glutathione and iron content. **B.** Intracellular glutathione content. **C.** Intracellular iron content. Asterisk (*) denotes significant difference. Letter a indicates significant differences against the WT strain growing in the control media. Error bars represent standard deviation (SD) values. In total, three biological replicates were performed (Mann–Whitney test, p < 0.05).

In this context, turning our focus to the homeostatic response, we quantified the intracellular levels of glutathione and iron in the WT and Δ*gsh* strains exposed to 0.5 mM FeCl_3_. As anticipated, a substantial reduction (two-times) in reduced glutathione content was evident in the Δ*gsh* strain compared to the WT counterpart (**Figure 2B**). Notably, iron excess significantly diminished glutathione levels in the WT strain, reinforcing the idea that this molecule is actively consumed to counteract metal toxicity. In the Δ*gsh* strain, glutathione levels showed an additional 30% reduction, yet this decline did not impact bacterial viability, suggesting the activation of complementary homeostatic pathways.

To explore this compensatory response further, we measure the intracellular content of iron of each strain (**Figure 2C**). Under basal growth conditions, no significant differences surfaced between WT and Δ*gsh* strains. However, under iron exposure, the Δ*gsh* strain exhibited a remarkable threefold reduction in intracellular iron levels compared to the WT (**Figure 2C**). This observation highlights a compensatory metal homeostatic strategy used by *E. faecalis*, whereby the bacterium reduces its iron content to cope with the reduced amount of glutathione produced in the absence of γGCS-GS.

Additionally, considering the importance of glutathione and iron for the bacterium, our findings underscore a potential interplay between glutathione synthesis and iron homeostasis in *E. faecalis*, shedding light on potential metabolic adjustments on a global level.

### 3. *E. faecalis* specifically reconfigures its global transcriptional response in absence of γGCS-GS

To determine whether the decrease in intracellular glutathione concentration affects the global transcriptional response, we performed a gene expression assay comparing the WT strain against the mutant Δ*gsh* strain. The analysis revealed a significant alteration in the transcriptional landscape when comparing the WT to the Δ*gsh* strain under basal growth (**Figure 3A**). Specifically, a total of 310 genes showed changes in the expression under absence of γGCS-GS enzyme. This strongly indicates that disruption of glutathione biosynthesis produces a broad transcriptional change compared to normal conditions with glutathione biosynthesis intact.

**Figure 3.**
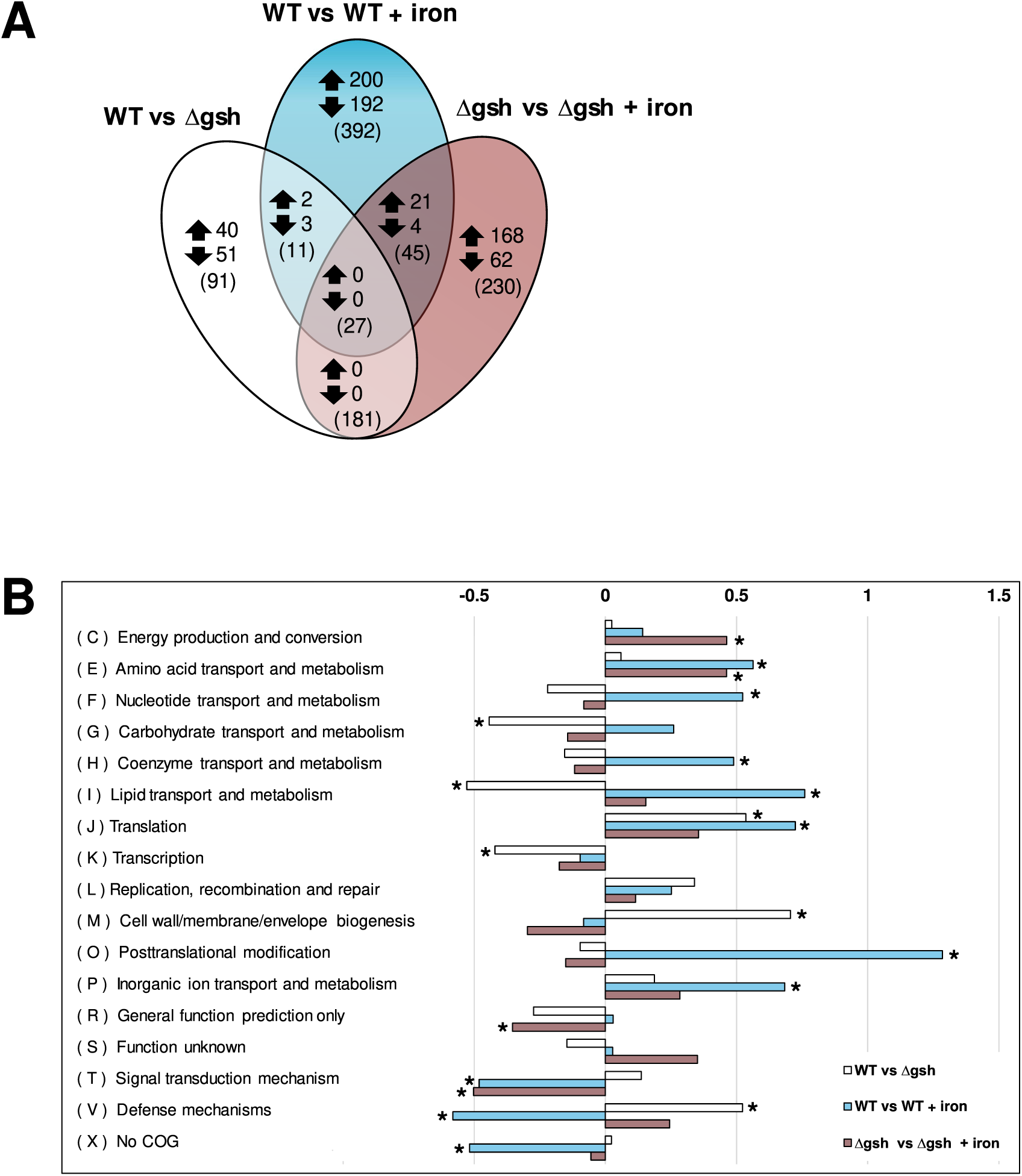
Comparative of transcriptional changes of *E. faecalis* WT and *Δgsh* exposed to iron. **A.** Venn diagram of sets of differentially expressed genes. The diagram declares the number of genes that were up-regulated (upward arrow) or down-regulated (downward arrow) shared by the comparisons. Numbers in parentheses indicate the total of genes differentially expressed. **B.** Enrichment analysis. The values are presented as Log(2) of the of total differential expressed genes normalized by the total genes in the genome grouped per COG category. Asterisk denotes significant enrichment (P < 0.05) compared to the total proportion in the bacterial genome.

Under iron excess (3 h, 0.5 mM FeCl_3_), WT strain exhibited differential expression in 475 genes, reinforcing its well-documented sensitivity to iron availability fluctuations (6). However, the most striking transcriptional reconfiguration was observed in Δ*gsh* strain, where 483 genes exhibited altered expression patterns, reflecting a substantial shift that differs from WT strain. Notably, only a small subset of genes (n=25) overlapped between the Δ*gsh* and WT strains under iron exposure, highlighting the distinct regulatory adjustments driven by glutathione depletion. One possible explanation for this response is that the basal gene expression profile of both strains differs even before iron exposure (**Figure 3A**), pre-conditioning the mutant strain to activate alternative regulatory mechanisms upon stress. In this regard, the decrease in glutathione content leads to a global transcriptional shift, which would also be impacting a decrease in the intracellular metal content (**Figure 2C**).

To gain further insights into the metabolic processes involved in this specific response, we performed a Clusters of Orthologous Groups (COG) enrichment analysis (**Figure 3B**). The reduction in glutathione content without iron treatment impacts mainly by decreasing the representation of genes linked to basal metabolism (carbohydrates (n=12) and lipids (n=4)) while increasing the biogenesis of the membrane and peptide glycan (n=18).

After treatment with iron, the reduction in glutathione content generates a global metabolic change similar to the WT, where processes linked to basal metabolism, such as nucleotide/carbohydrate biosynthesis and post-translational modifications, are significantly increased in both the WT and Δ*gsh* strains. These results suggest that glutathione depletion does not completely disrupt the metabolic adaptation to iron stress but may instead redirect resources, likely toward pathways controlling the toxic effects of iron.

Notably, category C (energy production and conversion) was significantly overrepresented only in the glutathione mutant under iron exposure. This suggests that Δ*gsh* strain may require additional ATP or reducing power to sustain antioxidant enzyme activity, highlighting a potential metabolic cost associated with compensating for glutathione deficiency.

In terms of transcriptional regulation, the decrease of glutathione leads to a decrease in the transcriptional repressor ArgR (EF0983, regulator of arginine metabolism) (25, 26), and an increase in the expression levels of global transcription factors belonging to the LysR families (EF0644, EF0923 and EF2958) and TetR (EF0791, EF0787, EF2203, EF2066), which are involved in basal metabolism, energy generation and general stress responses, therefore suggesting that glutathione depletion induces regulatory adjustments at the transcriptional level which could lead to important changes in the basal metabolism of *E. faecalis*.

### 4. Low levels of glutathione impacts over the amino acid concentrations

Considering the gene expression results, with the aim of identifying which metabolites could be affected by the decrease in glutathione levels, a metabolomic approach was carried out to determine the compounds involved in the response to iron exposure in both the WT and mutant strain (**Table 1**).

**Table 1.**
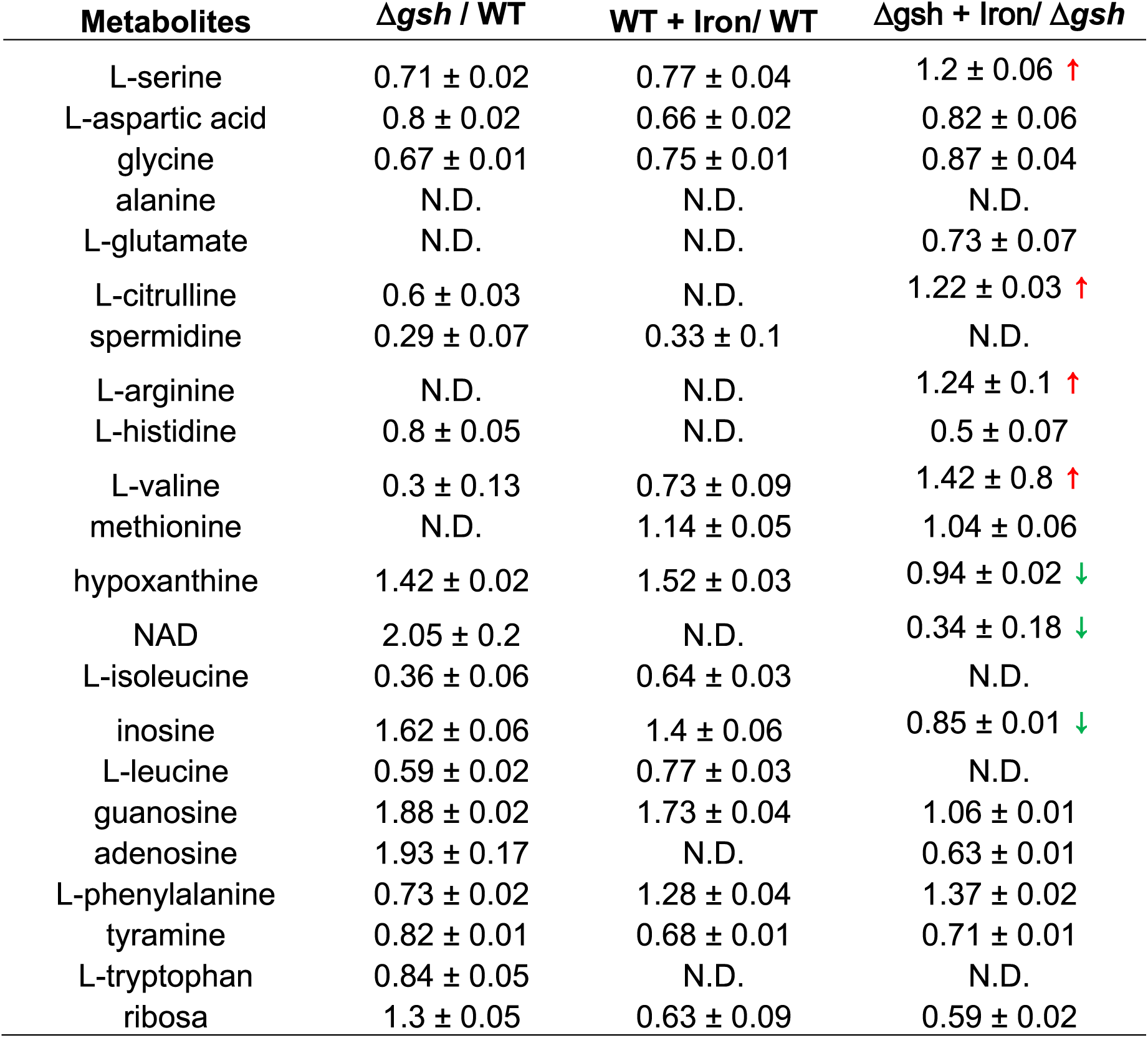
Relative abundance of metabolites in *E. faecalis* WT and *Δgsh* exposed to iron. The values represent the average of the rate of change between both conditions of 3 independent replicates. A t-test was used to evaluate statistical differences between means from the two conditions (N.D. no significant differences, p < 0.05). Red and green arrows represent an increase or decrease of the metabolite in the mutants in relation to the WT exposed to iron, respectively.

In general terms, the decrease in glutathione levels significantly impacts the basal metabolism of the bacteria. Significant changes were observed in nearly all quantified amino acids, as well as in NAD synthesis, nucleotide, and polyamine metabolism.

Under metal exposure, consistent with the transcriptional level findings, significant changes were detected in the concentration of several key metabolites in the Δ*gsh* strain treated with iron. There was an increase in the concentration of charged amino acids such as L-arginine and L-citruline (metabolism of arginine), polar amino acids such as L-serine, and the apolar residue L-valine. Notably, arginine metabolism can serve as a sole source of nitrogen, carbon and energy (27). For example, the deimination of this amino acid allows the generation of ATP from ADP and phosphate while releasing ammonia (28). This increase in arginine levels in the mutant exposed to iron directly correlates with the decrease in the abundance of the repressor ArgR, which could be regulating the *arcA* genes (EF0104, encoding arginine deiminase), *arcF* (EF0105, encoding ornithine carbamoyltransferase) and *argC* (encoding carbamate kinase), all of which are derepressed in the mutant compared to the WT.

The increase in arginine observed during the decrease in glutathione content could also be correlated with the detected increase in polyamines. Arginine and its precursor, ornithine, serve as substrates for synthesizing putrescine and spermidine, two major polyamines widely involved in ROS control, energy production and protein synthesis (29–31). Conversely, there is a reduction in the concentration of metabolites related to ATP catabolism (inosine and hypoxanthine) and NAD in the mutant strain. The decrease in these compounds suggests greater utilization of these metabolites, which directly correlates with global gene expression assays, supporting the increased overrepresentation of genes involved in energy production and conversion mechanisms.

### 5. Compensation pathways are related to arginine metabolism

Currently available is the Genome-scale metabolic model (GSMM) for *E. faecalis* published in 2015 (32), which was recently expanded and curated (33). The metabolic model of *E. faecalis* consists of 1,336 reactions and 1,293 metabolites, including all listed in **Table 1**. Regarding the impact of the absence of the γGCS-GS enzyme under basal condition (Sup. Figure 3), there is an envelope expression of genes linked to purine and pyrimidine synthesis pathways, which directly correlating with the increase in metabolites associated with nucleic acids. This is coupled with the repression of genes associated with amino acid synthesis pathways, such as L-phenylalanine and L-tryptophan, explaining the decrease in these amino acids as a possible substitute for the decrease in intracellular glutathione content.

In response to metal treatment, **Figure 4** shows the integration of transcriptomics and metabolomics data within the metabolic model for both the WT and mutant strains exposed to iron. In both strains, metal exposure leads to the activation of metabolic pathways linked to arginine metabolism. In the WT strain, routes beginning mainly from the urea cycle which are conducive to the generation of basal metabolites of the nucleotide type (pyrimidines) and amino acids of the alanine, aspartate and glutamate types, the latter used in the synthesis of glutathione, added to the activation of routes associated with the basal metabolism of carbohydrates and fatty acids. In contrast, metabolism in the mutant strain derives through arginine synthesis pathways, which, as previously mentioned, seem to lead to the production of metabolites related to energy generation.

**Figure 4.**
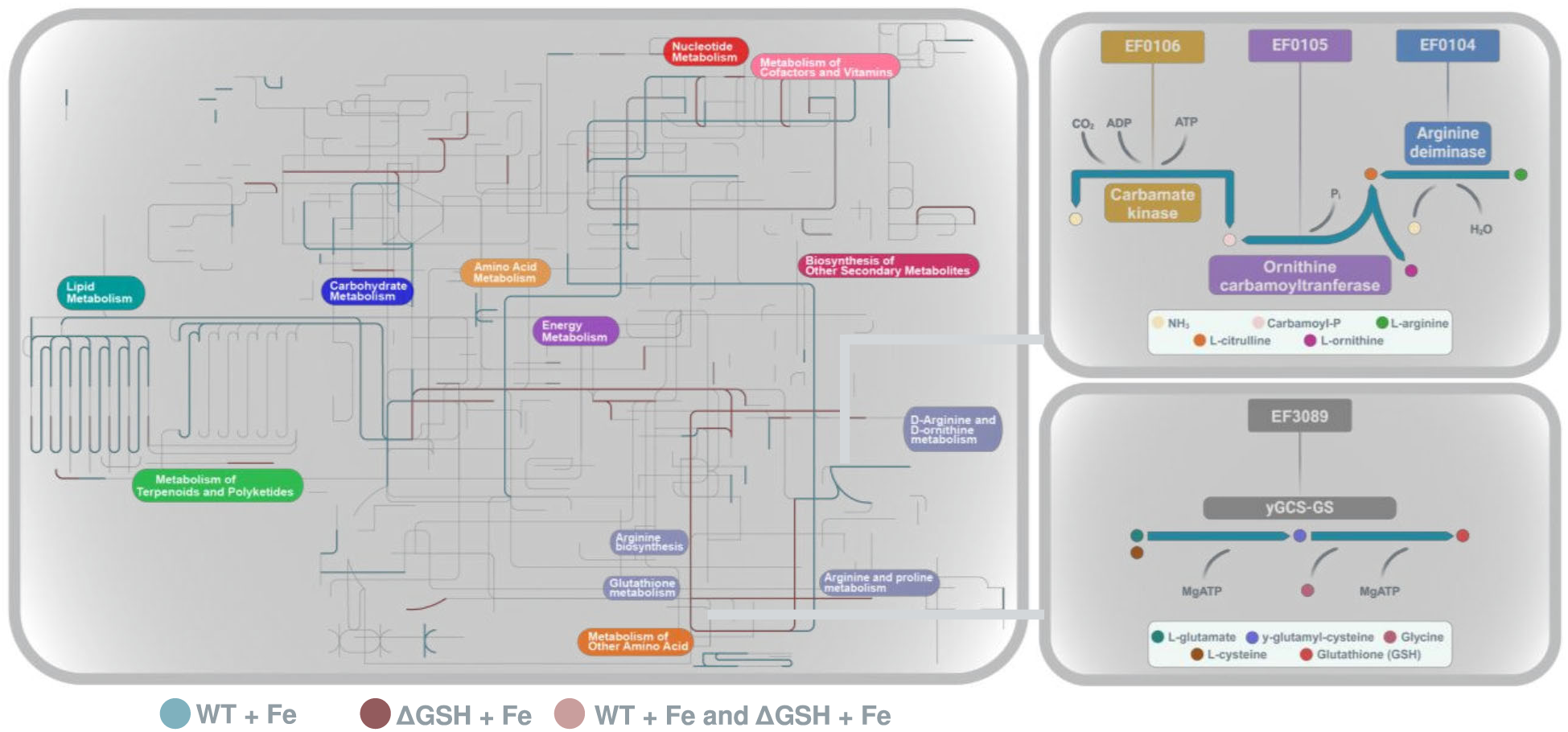
Metabolic map of *E. faecalis* according to gene expression levels. A pictorial representation of the fluxes involved in the response of *E. faecalis* WT and Δ*gsh* exposed to iron. Each reaction was marked based on the enzyme encoded by the differentially expressed gene from each condition. The integrated representation was generated using Ipath 3.0.

According to these assumptions, arginine metabolism appears to act as a compensatory mechanism for reduced glutathione levels. To explore this, an *in silico* double mutant (Δ*gsh*Δ*arc*) was generated by removing genes related to arginine catabolism (*arc* operon; EF0104, EF0105 and EF0106) and glutathione synthesis (EF3089) from the metabolic model.

Using flux balance analysis (FBA) along with parsimonious criteria (pFBA), the optimal flux distributions for all model reactions and biomass production were calculated for the WT and double mutant models. Overall, there were no differences in biomass production between the two models (BM = 0.35 mmol / (gDW x h)), indicating the robustness of this bacterium’s metabolism to compensate for the elimination of specific metabolic pathways, including those associated with amino acid synthesis.

Among the changes, four pathways linked to glycerol metabolism were highly affected (**Figure 5**). Conversely, the glycerol reductive pathway was highly favored. First, there was an increase in the unidirectional flux of glycerol-3-phosphate synthesis, coupled with a decrease in the consumption of this metabolite, favoring the generation of dihydroxyacetone phosphate, which is consumed through glycolysis to form pyruvate and subsequent energy production.

**Figure 5.**
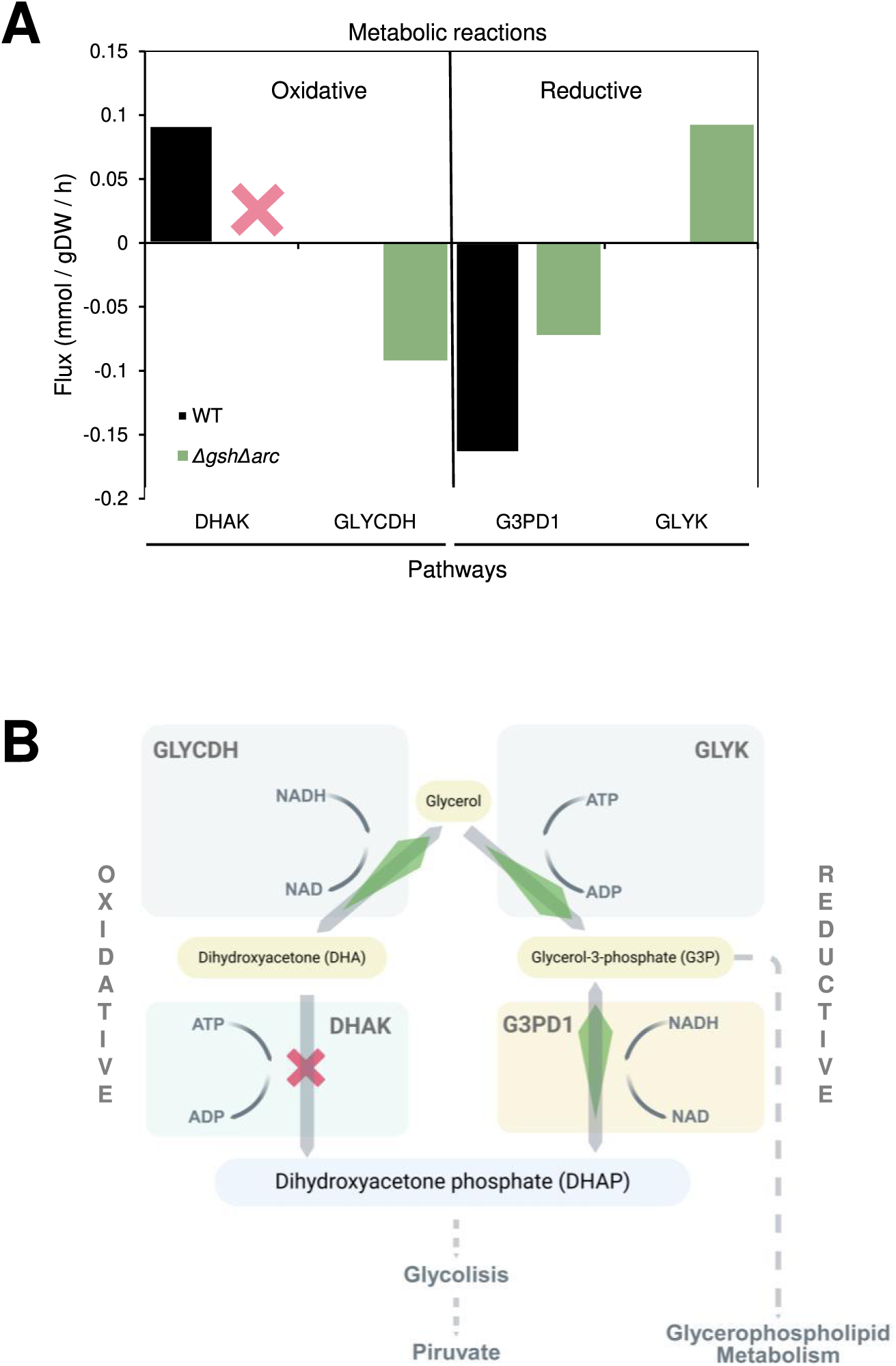
FBA results and schematic overview of compensatory fluxes in the double mutant metabolic model. The effects of the double-mutations were generated by removing the reactions directly from the metabolic model corresponding to each enzyme encoded by the genes *gsh* gene (γGCS-GS enzyme) and *arc* operon (anerobic arginine catabolism). **A.** Bars corresponds to reaction fluxes of glycerol metabolism. **B.** Schematic overview of glycerol metabolic fluxes. An increase in the thickness of the green arrows and the X red letter indicates the directionality or the blockage of the reactions for the fluxes that changed in the double mutant, respectively.

The activation of both arginine and glycerol metabolism suggests the existence of compensatory mechanisms in *E. faecalis* capable of adjusting bacterial metabolism to maintain cell viability, even under stress conditions such as iron exposure.

## DISCUSSION

The equilibrium between oxidants and reductants is crucial for the homeostasis of an organism. In this sense, the iron, through the Fenton-Haber Weiss reaction, amplifies oxygen toxicity by producing ROS (34). It is believed that tightly controlling iron metabolism, coupled with regulating defenses against oxidative stress, is essential in mitigating its harmful effects (35). In this context, *E. faecalis* has emerged as a prominent bacterial model for investigating transcriptional and metabolic adaptations in response to iron stress, offering valuable insights into the regulatory networks that coordinate metal homeostasis and oxidative stress responses (6, 15).

Here, we provide new evidence on how glutathione depletion reshapes the global transcriptional and metabolic responses of *E. faecalis* under iron excess. Prior research has established the essential role of glutathione in conferring tolerance to oxidative stress (10, 36, 37) and in mitigating the toxic effects of divalent metal ions (37). Our findings reveal that while glutathione deficiency does not impair bacterial viability, it triggers broad transcriptional and metabolic reprogramming, likely as a compensatory strategy.

### Bifunctionality of γGCS-GS enzyme in *E. faecalis*

According to our *in silico* and metabolic results and supporting previous analyses (20), *E. faecalis* utilizes a bifunctional γGCS-GS enzyme for glutathione biosynthesis, rather than the two-step pathway present in other bacteria and eukaryotes (38). This characteristic, also observed in *S. agalactiae* and *L. monocytogenes*, suggests an evolutionary adaptation that enhances glutathione synthesis efficiency.

Notably, γGCS-GS activity in *E. faecalis* is not inhibited by glutathione levels, as observed in *S. agalactiae*, unlike other bacteria in where it exists in monofunctional variants for glutathione synthesis (20, 39). The lack of feedback inhibition may confer an advantage under oxidative stress conditions, allowing continuous glutathione production. In this sense, the bifunctionality of the enzyme, coupled with its lack of product inhibition, generates a more efficient process in the generation of glutathione, which could be essential in optimizing the antioxidant capacity of pathogenic organisms such as *E faecalis*, *Listeria monocytogenes* or *S. agalactiae* (40, 41) to cope more effectively with the oxidative stress generated by the host.

### *E. faecalis Δgsh* strain characterization under iron excess

The deletion of γGCS-GS was not lethal for *E. faecalis*, consistent with observations in other bacteria such as *Pseudomonas aeruginosa* and *Streptococcus pneumoniae* mutants for glutathione biosynthesis (36, 37). In addition, the glutathione levels in the Δ*gsh* strain were significantly lower than in the WT, likely due to compensation through glutathione uptake and regeneration pathways (42, 43). However, these mechanisms appear less efficient than the de novo biosynthesis catalyzed by γGCS-GS, hence the presence of intracellular glutathione is anticipated.

Glutathione levels decreased under iron exposure in both WT and Δ*gsh* strains, consistent with an oxidative stress response. Similar conditions (0.5 mM iron) have been shown to induce oxidative stress in *E. faecalis*, characterized by reduced glutathione levels and upregulation of antioxidant genes such as *sodA*, *katA*, *trx*, and *msrA* (15). Comparable oxidative stress responses have been observed in other bacteria, including *A. baumannii* and *E. coli* (44, 45).

In our case, we found similar results during iron increase in the WT strain, showing a reduction in glutathione levels and upregulation of genes for thioredoxin reductase (EF1338) and peptide methionine sulfoxide reductase (EF1681). Therefore we can suggest that *E. faecalis* faces a possible scenario of oxidative stress under iron increase. In contrast, the Δ*gsh* strain with iron increase did not exhibit the expression changes observed in the WT, despite the reduction of glutathione levels. We attributed this to a compensation scenery that appears to be dedicated to avoid an increase in intracellular iron levels, as indicated by the significantly reduced levels of intracellular iron in the Δ*gsh* strain compared to the WT.

In *P. aeruginosa*, glutathione mutants reduce iron uptake by decreasing siderophore production (36). Although the Δ*gsh* strain of *E. faecalis* did not show significant changes in iron uptake gene expression (6, 15), its reduced intracellular iron levels suggest compensatory mechanisms linked to protein efficiency rather than abundance. Additionally, glutathione may act as an intracellular iron buffer or retainer of Fe-S cofactors (46, 47). This could further support the observed phenotype in the mutant.

### Transcriptional reconfiguration of *E. faecalis* Δ*gsh* strain

Our analysis of the global transcriptional changes under glutathione depletion in *E. faecalis* revealed a broad transcriptional shift in the Δ*gsh* strain. Similar features have been reported in other microbial species. For example, a microarray analysis in *S. cerevisiae* showed that a mutant strain for γ-glutamylcysteine synthetase (GSH1), encoding the rate-limiting enzyme that participates in the first step of glutathione biosynthesis, presented differential expression of 189 genes under glutathione depletion (48). Similarly, study in the glutathione auxotrophic *Streptococcus pyogenes*, revealed an extensive transcriptional perturbation under glutathione decrease condition (11). Specifically, total RNA-sequencing (RNA-seq) of mutant strain for a key player in glutathione import in *S. pyogenes* (HKU16Δ*gshT* strain), showed expression changes in 165 genes. Interestingly, the majority of these genes (75%) were downregulated, including several virulence-determinant genes.

Given this context, the different transcriptional response that we observed between *E. faecalis* WT and Δ*gsh* was expected. The Δ*gsh* strain exhibited reduced expression of genes related to basal metabolism, including nucleotide and carbohydrate biosynthesis, as well as post-translational modifications. This shift may reflect an adaptive strategy to limit metabolic activity when glutathione is depleted, reducing ROS production (49). Similar responses have been observed in *A. baumannii* and glutathione-auxotrophic *S. pyogenes*, where metabolic downregulation occurs under oxidative stress (11, 50). Notably, the reduction in basal metabolism in Δ*gsh* was evident even under non-stress conditions (**Figure 3**), reinforcing the idea of a systemic adaptation to glutathione deficiency.

The transcriptional shift observed in *E. faecalis* implies the participation of transcriptional regulators, with an example of this scenario reported in *P. aeruginosa*. The redox balance performed by glutathione upregulates T3SS gene expression via the global transcription factor Vfr, which senses glutathione and activates T3SS expression (51).

To our knowledge, *E. faecalis* does not present reports of transcription factors that sense glutathione levels; however, in our analysis, the decrease in glutathione resulted in the downregulation of transcriptional repressor ArgR and the upregulation of transcription factors from the LysR and TetR families involved in basal metabolism, energy generation and general stress. This suggests that transcriptional control mechanisms affected by glutathione levels are present in *E. faecalis*. A previous study introduced a global transcriptional regulatory model for *E. faecalis* (14). Interestingly, the operon *arc* (EF0104, EF0105 and EF0106), contain in their promoter region a DNA binding site for the regulator ArgR, which is consistent with the transcriptional changes of these genes in the mutant under the iron exposure.

### Compensatory metabolic changes in *E. faecalis*

Our metabolic analysis revealed that under iron stress, the Δ*gsh* strain redirects its metabolic flux toward arginine synthesis pathways, apparently leading to energy generation and reductive power production. Thus, in a scenario where *E. faecalis* is faced with increased iron under glutathione deficiency, one way to mitigate the harmful effects of iron is to reduce intracellular levels of the metal. This metabolic shift may enhance energy production and support ROS detoxification. Consistently, we observed an increase in L-arginine levels and the derepression of genes involved in its synthesis, including reduced expression of the repressor *ArgR*, highlighting the importance of the metabolic pathways involved, which could act as a compensatory system when glutathione levels are compromised.

Unlike amino acids such as histidine, isoleucine, methionine and tryptophan, which cannot be synthesized by *E. faecalis* (essential amino acids) (52), arginine synthesis pathways play a fundamental role in bacterial metabolism. The changes in metabolic fluxes in the mutant significantly impacted arginine synthesis and pathways related to NAD metabolism, indicating a redistribution towards stress protection in response to decreased glutathione content. This phenotype likely does not affect the overall biomass of the bacteria, as supported by viability assays, where no changes were observed in the growth of Δ*gsh* bacteria during metal exposure compared to the WT strain.

The derepression of the *arc* operon in the mutant strain under iron stress suggests that anaerobic arginine catabolism acts as a compensatory mechanism for glutathione deficiency. Arginine metabolism has been linked to oxidative stress resistance in several bacteria, including *Salmonella*, where it protects against ROS-induced damage (53, 54). We hypothesize that L-arginine metabolism is part of the *E. faecalis* response to alleviate the ROS produced under conditions of increased iron and decreased glutathione.

Through the analysis of fluxes in the double mutant model, we propose glycerol metabolism as a possible compensatory pathway activated in response to the absence of arginine synthesis pathways linked to genes responsive to glutathione depletion. The pathway for glycerol catabolism involves a dismutation process. The deletion of the DHAK pathway leads to an increase in dihydroxyacetone, which is metabolized towards glycerol via the GLYCDH pathway, showing an increase in flux in the same direction. This reverse flux essentially leads to the coupled reductive pathway, where glycerol is converted to 3-hydroxypropionaldehyde and further to 1,3-propanediol (both pathways exhibit increased fluxes in the double mutant). It is important to note that even in the double mutant, the bacterial biomass remains unaffected, indicating that the metabolic system in *E. faecalis* remains robust even in the absence of arginine and glutathione synthesis pathways.

Considering that *E. faecalis* is a facultative aerobic fermentative bacterium, its metabolism tends to ferment carbohydrates to produce energy, generating lactic acid as one of the final products through reductive reactions. This aligns with the results obtained in the model, reflected in the involvement of the *arc* operon and the activation of the glycerol reductive pathway as compensatory mechanisms.

In summary, these results demonstrate the existence of compensatory mechanisms in *E. faecalis*, which are directly involved in metal homeostasis (reduction of iron content) and others associated with transcriptional changes that directly correlate with increases or decreases in metabolic fluxes, primarily aimed at generating greater energy through the use of reductive pathways. All mechanisms aim to cope with the decrease in glutathione content in response to an oxidative agent such as iron.

## METHODS AND PROTOCOLS

### *In silico* analysis of *E. faecalis* γGCS-GS protein

Glutathione synthetase (GS) molecular 3D model was generated by SWISS-MODEL program (55), using information from the crystalline structure of γGCS-GS from *Streptococcus agalactiae* as a template. The final model was displayed with VMD v1.9.4 software (56). Glutathione protein homologous in *Lactobacillales* order organisms were recovered by BlastP on the National Center for Biotechnology Information (NCBI) website. The template gene was *E. faecalis gshF* gene EF3089. Global glutathione alignments were performed using ClustalW as described in previous work (6).

### Strains, growth conditions and iron treatments

All strains (*E. faecalis* OG1RF and *Δgsh*) were grown in N medium (Peptone 1%, yeast extract 0.5%, Na_2_HPO_4_ 1% and glucose 1%) (57). Mutant glutathione strain (*Δgsh*) of *E. faecalis* OG1RF were constructed using the pTEX4577 vector system (57, 58). For all the experiments, both strains were independently cultured overnight in N medium broth at 37°C and in triplicate. The next day, cells were diluted in two parallel cultures (control and iron) to a final OD_600nm_ of 0.05 and then grown at 37 °C and 160 rpm. The iron culture was supplemented with concentration of 0.5 mM as previously determinate as a nonlethal (6), 1, 2 and 4 mM of FeCl_3_. Bacterial growth was monitored each hour over 8 h by OD_600nm_. Transcriptomic and metabolomics samples were obtained from WT and *Δgsh* strains after 3 h with 0.5 mM of FeCl_3_ and without iron added (control).

### Intracellular iron and glutathione content

After the iron treatments (control or excess), the intracellular metal and reduced gluthatione content for each culture was determined as described below (57). Both strains growing in the different iron conditions and at the same growth stage (mid-exponential, 3 h of growth cultures according to the growth curves), were centrifuged and then suspended in 200 μl of HNO3 (Merck) and digested during 24 h at 65°C. After the acid lysis, total metal composition was determinate by total reflection X-ray fluorescence (TXRF). The data were expressed as µg/L per liter, normalized per total cell culture (absorbance). As previously described (59), the content of reduced glutathione (GSH) was quantified using the ortho-phthalaldehyde (oPT) method, The results are presented as s nmol GSH/mg protein. Values represent the average data of three measurements for independent biological replicates. The statistical analysis was done using an Mann–Whitney test, p < 0.05.

The glutathione content was determined as described previously (60). Each sample was suspended in 1 mL of buffer containing 50 mM Tris–HCl (pH 8.0), 5 mM EDTA, 0.1% SDS, and 0.1 mM DTNB (5,5’-dithiobis-(2-nitrobenzoic acid)). The suspension was incubated at 37 °C for 30 minutes, homogenized by vortexing for 10 seconds, and centrifuged at 14,000×g for 10 minutes. The absorbance of the supernatant was measured spectrophotometrically at 412 nm. Glutathione levels were determined using the molar extinction coefficient of oxidized DTNB (1.36×10⁴ M⁻¹ cm⁻¹). Calibration curves were constructed using GSH solutions of known concentration.

### Global expression assays

Gene expression analysis was performed using a chip of four arrays of 72 K (catalog number A7980-00-01) from NimbleGen Systems, Inc. Each array contains 72,000 oligonucleotides 60-mer, with an average of 11 oligos designed for the 3,114 open reading frames of *E. faecalis* OG1RF that represent twice the genome of the bacterium (two technical replicates per arrays). Next, for WT and *Δgsh* strain four independent hybridizations (two biological replicates per treatment paired with their respective controls) were performed by the manufacturer (Nimblegen) on equal conditions in a single glass, thus reducing variability between hybridizations (pre-hybridization, hybridization and washing steps). Scanning of the slides and data analysis also were carried out by Nimblegen. The gene expression from Fe-treated WT cells were compared to that of untreated cells; data from Fe-treated *Δgsh* mutant cells were compared to untreated *Δgsh* cells and data from *Δgsh* strain were compared to WT cells. Student’s t-test statistics were used to identify significantly different gene expression levels with a P < 0.05 and a 4-fold magnitude of change between the average value of each gene and its corresponding reference, using the DNASTAR software Array star 3.0, such as the previous report (57). Data available in GEO database (ID GSE34432 and GSE275402). For the gene enrichment analysis, the complete set of genes of *E. faecalis* OG1RF was used on BLASTP analysis against the Cluster of Orthologous Groups database (COG). Based on the retrieved annotations, an enrichment analysis of COG categories was carried out for the gene sets of *E. faecalis* OG1RF using Fisher’s exact test with Benjamini– Hochberg multiple testing correction. Categories with corrected p-value <0.05 were considered enriched.

### Metabolomic data

The sampling, quenching, and intracellular metabolite extraction will be done based on reported protocols (61). Briefly, 30 milliliters of culture broth was sampled in triplicate and quenched with cold glycerol-saline solution (3:2), then centrifuged at −20°C. The cell pellets were resuspended in cold glycerol-saline solution (1:1) and centrifuged again at −20°C. Intracellular metabolites were extracted from the cell pellets using cold methanol-water solution (1:1) at −20°C, followed by three freeze-thaw cycles. After centrifugation at −20°C, the supernatant was collected. The cell pellet was then resuspended in cold pure methanol (−20°C) for a second extraction, centrifuged at −20°C, and the supernatant was pooled with the previous one. The cell pellet was resuspended in bi-distilled water, centrifuged, and the supernatant collected. Finally, twenty milliliters of cold bi-distilled water (4°C) was added to the metabolite extracts, which were then frozen and freeze-dried. Next, metabolites detection and was realized by capillary electrophoresis (CE) coupled with electrospray ionization time-of-flight mass spectrometry (ESI-TOF-MS). Normalization and quantification of metabolites were performed following previous validated analytic methods (62). To identify significant differences between mean values between conditions, a two-tail t-test was performed assuming independence of groups and equal variances. The null hypothesis was rejected if the p-value was less than 0.05.

### *E. faecalis* Genome Scale Metabolic Model (GSMM) analyses

For metabolic network construction, differentially expressed genes and selected metabolites were used to analyze the putative metabolic output through the genome scale metabolic model available for *E. faecalis* (32, 33). Three *in silico* mutants were designed using this model. First, the GSH mutant was generated by deleting the reactions associated with the gene EF3089 (*gshF*), corresponding to the reactions GLUCYSL and GTHS. For the arginine mutant, the reactions linked to genes EF0104-0106, specifically *argd*, *ocbt*, and *ck* were removed. Lastly, a double mutant was created by eliminating all reactions associated with both EF3089 and EF104-106 genes. For all these mutants, the maximum growth rate was defined as the objective function, and flux balance analysis (FBA) along with parsimonious flux balance analysis (pFBA) was performed to calculate the optimal flux distributions across the metabolic network. Specific uptake rates were used as constraints, aligning the flux through the biomass equation with the specific growth rate (mmol gDW^−1^ h^−1^). This allowed FBA to determine optimal yield strategies, given that the maximum growth rate depends on the input fluxes (63, 64). Both FBA and pFBA were implemented using the COBRA Toolbox in MATLAB, with GLPK employed to solve the optimization problems. iPath 3.0 was used to visualize the metabolic pathways of the GSMM (65). Pathways and specific reactions were highlighted based on changes in gene expression between the Δ*gsh* and WT strains, with and without iron exposure (*E. faecalis* NCBI ID 226185).

## ACKNOWLEDGEMENTS

This work was supported by Proyecto ANILLO regular ANID ACT210004, Center for Mathematical Modeling, Apoyo a Centros de Excelencia ACE210010; ANID Millennium Science Initiative Program ICN2021_044; Fondo Basal FB210005; ANID FONDECYT 1190742, 1230194, 1230195 and 1230724, Fondo Interdisciplinario y Proyecto Núcleo UOH; FONDAP 1513001; Proyecto Postdoctoral ANID 3220080; Universidad de Chile: grant Apoyo a la Infraestructura para la Investigación INFRA037/2023.

## SUPPLEMENTARY INFORMATION

Document S1. Figures S1–S3.

